# Structure-Function Analysis of the FCRL5-IgG1 Fc Complex Reveals an Unappreciated Effect of Fc-Silent Antibodies on B cells

**DOI:** 10.64898/2026.04.16.718972

**Authors:** Bart M. Herpers, Mo Guo, Sanghwan Ko, George Delidakis, Jin Eyun Kim, Chang-Han Lee, Mohamed I. Gadallah, Jennifer S. Brodbelt, Y. Jessie Zhang, George Georgiou

**Author notes:** Corresponding Authors: Y. Jessie Zhang and George Georgiou.

## Abstract

Human Fc receptor-like 5 (FCRL5) is a low-affinity IgG Fc receptor expressed on various B cell subsets and a potential therapeutic target. We discovered that commonly used Fc-silencing mutations, designed to prevent interactions between the Fcγ receptors on immune cells and the Fc domain of therapeutic IgG, do not prevent binding to FCRL5. As a result, unintended interactions between Fc-silent therapeutic IgG and human B cells may occur. We isolated a well-expressed variant of the Fc-binding portion of human FCRL5 by directed evolution and used structural modeling to guide the engineering of a human IgG1 Fc variant with approximately 100-fold higher affinity for FCRL5, enabling us to produce FCRL5:Fc complexes in solution. Native mass spectrometry, size exclusion chromatography, and the crystal structure of the FCRL5- IgG1 Fc complex solved at 3.4 Å indicate that the two proteins bind in a 1:1 stoichiometry. Furthermore, the structure revealed that FCRL5 binds to IgG1 Fc in a manner completely distinct from that of previously characterized Fc-binding proteins, such as Fcγ receptors, explaining why most Fc-silencing mutations do not disrupt FCRL5 binding. We demonstrate that selective cross-linking of FCRL5 with the B cell receptor (BCR) *in cis*, using Fc-engineered antibodies with either physiological or enhanced FCRL5 affinity, inhibits Ca^2+^ flux in FCRL5-expressing B cells. We compare this effect with the selective co-ligation of FcγRIIb with the BCR. Our work demonstrates that FCRL5 interacts with human IgG Fc in a distinctive manner and that engagement of FCRL5 by Fc-silent therapeutic IgG could influence B cell function.

## Introduction

The physiological and pathophysiological roles of IgG antibodies are critically dependent on a plethora of interactions mediated by the Fc domain. The IgG Fc domain in immune complexes interacts with effector type I Fc receptors, which are widely expressed on myeloid and lymphoid cells, as well as on some non-hematopoietic cells (*1*). This interaction elicits inflammatory responses via FcγRI, FcγRIIa, FcγRIIIa and FcγRIIIb, and anti-inflammatory effects through FcγRIIb (*2*, *3*). Additionally, the IgG Fc domain interacts with type II Fc receptors (*1*), C1q (*4*), FcRn (*5*), TRIM21 (*C*), and pathogen Fc receptors (*7*, *8*). For many therapeutic antibodies, particularly those that function mainly through target blockade or agonism, it is crucial to reduce, and ideally eliminate, binding to all FcγRs and to C1q. Various combinations of mutations that “silence” the Fc domain have been employed in numerous approved or clinical-stage therapeutic antibodies (*9*), with a commonly used combination being L234A, L235A, P329G (LALAPG) (*10*). However, while the existing Fc-silencing mutations have been established to eliminate or significantly attenuate antibody-dependent cell cytotoxicity (ADCC), phagocytosis (ADCP), and complement activation (CDC) (*11*), it is not known whether they may still have other subtle effects on adaptive or innate immune cells arising from hitherto unknown interactions.

In 2012, Colonna and colleagues demonstrated that IgG antibodies also bind to Fc receptor-like 5 (FCRL5), one of the five FCRL molecules encoded in tandem over approximately a 300 kb region on chromosome 1q21–22, telomeric to FCGR1 (*12*, *13*). The FCRL5 protein has nine extracellular immunoglobulin-like domains (D1-D9), of which the first three, D1-D3, are key for IgG binding (*12*, *14*). FCRL5 can bind to all four human IgG subclasses with a dissociation constant (*K*_D_) in the 1-10 μM range (*14*). Removal of the Fab regions has been reported to weaken the FCRL5-IgG interaction, with *K*_D_ values decreasing to 46-86 μM for IgG Fc (*14*, *15*). Separately, removal of the N297-linked IgG Fc glycan substantially weakens the interaction between IgG and FCRL5 (*14*).

FCRL5 expression is restricted to B cells (*16, 17)*. It is expressed on naïve and memory B cells within the tonsils and spleen of healthy individuals and on nearly all plasma cells, including long-lived bone marrow plasma cells (*18*). The cytoplasmic tail of FCRL5 encodes two canonical ITIM motifs and one ITAM-like motif (*16*). The exact biological function of FCRL5 remains unclear, with conflicting reports on whether its cross-linking activates or inhibits B cells (*19*–*21*). Nevertheless, a consistent finding across these studies is that FCRL5 inhibits B cell receptor (BCR)-mediated calcium mobilization when co-ligated with the BCR *in vitro*, likely through ITIM-mediated recruitment of the tyrosine phosphatase SHP-1 (*19*–*21*).

FCRL5 has been implicated in a variety of autoimmune diseases, including rheumatoid arthritis (RA), myasthenia gravis (MG), and systemic lupus erythematosus (SLE), among others (*22*). FCRL5 has also emerged as an important therapeutic target for B cell malignancies. High expression of FCRL5 occurs in multiple myeloma (MM) with near 100% penetrance (*23*, *24*), and a recent study suggests that it is also an important surface marker for MM-initiating cells (*25*). Despite emerging evidence on the biomedical significance of FCRL5, there is currently a paucity of information on its structure, function, and the molecular basis of IgG recognition. The low affinity of FCRL5 for human IgG Fc, coupled with the low expression yield of both the holoreceptor and the functionally significant D1-D3 domains, has hindered the structural analysis of the FCRL5-Fc complex. To overcome these limitations, we used directed evolution to develop an FCRL5 D1-D3 variant with enhanced expression in Expi293F cells. Additionally, we employed structural modeling to design an IgG1 Fc variant with approximately 100-fold lower *K*_D_ for FCRL5. We then determined the crystal structure of the FCRL5-IgG1 Fc complex at 3.4 Å resolution, which revealed that FCRL5 binds to IgG Fc in a manner completely distinct from that of all other known Fc-binding proteins, namely effector FcγRs, C1q, FcRn, and TRIM21. Importantly, we show that widely clinically used mutations that abrogate Fc effector functions (Fc-silencing mutations) have no effect on antibody binding to FCRL5. We demonstrate that co-ligation of FCRL5 with the BCR by antibodies having the clinically important LALAPG Fc-silencing mutations decreases Ca^2+^ flux in FCRL5-expressing B cells. This effect is independent of and distinct from FcγRIIb engagement, and mutations that enhance FCRL5 affinity further reduce Ca^2+^ flux. Our results shed light on the structural basis of the FCRL5:IgG interaction and highlight a possible liability of Fc-silencing mutations in therapeutic antibodies.

## Results

### Fc-silenced trastuzumab variants and IgG1 Fc hexamers bind to FCRL5

FCRL5 D1–D3 share overall sequence homology with the ectodomains of Type I FcγRs (FcγRI, FcγRII, FcγRIII); however, the amino acid residues that are key for FcγR:IgG binding are not conserved in FCRL5 D1–D3 (*26*). Prompted by this analysis, we examined whether Fc-silencing mutations designed to ablate interactions with the effector FcγRs are able to bind to the FCRL5 ectodomain (residues Q16-G851). Trastuzumab variants containing the clinically relevant Fc-silencing mutations LALAPG, E233P/L234V/L235A/G236del/S267K (named FcKO, (*27*)), or L234A/L235A/G237A/P238S/H268A/A330S/P331S (IgG1σ, (*28*)) had no binding to the inhibitory receptor FcγRIIb, whereas binding to FCRL5 was preserved (Figures 1A and 1B, S1A and S1B). Only the removal of the N297 Fc glycan through mutation of the critical T299 in the N-X-S/T motif impaired binding to the FCRL5 ectodomain, consistent with earlier reports (*14*). These results suggest that the FCRL5 binding site on IgG1 Fc is markedly different from that of the effector FcγRs.

**Figure 1.**
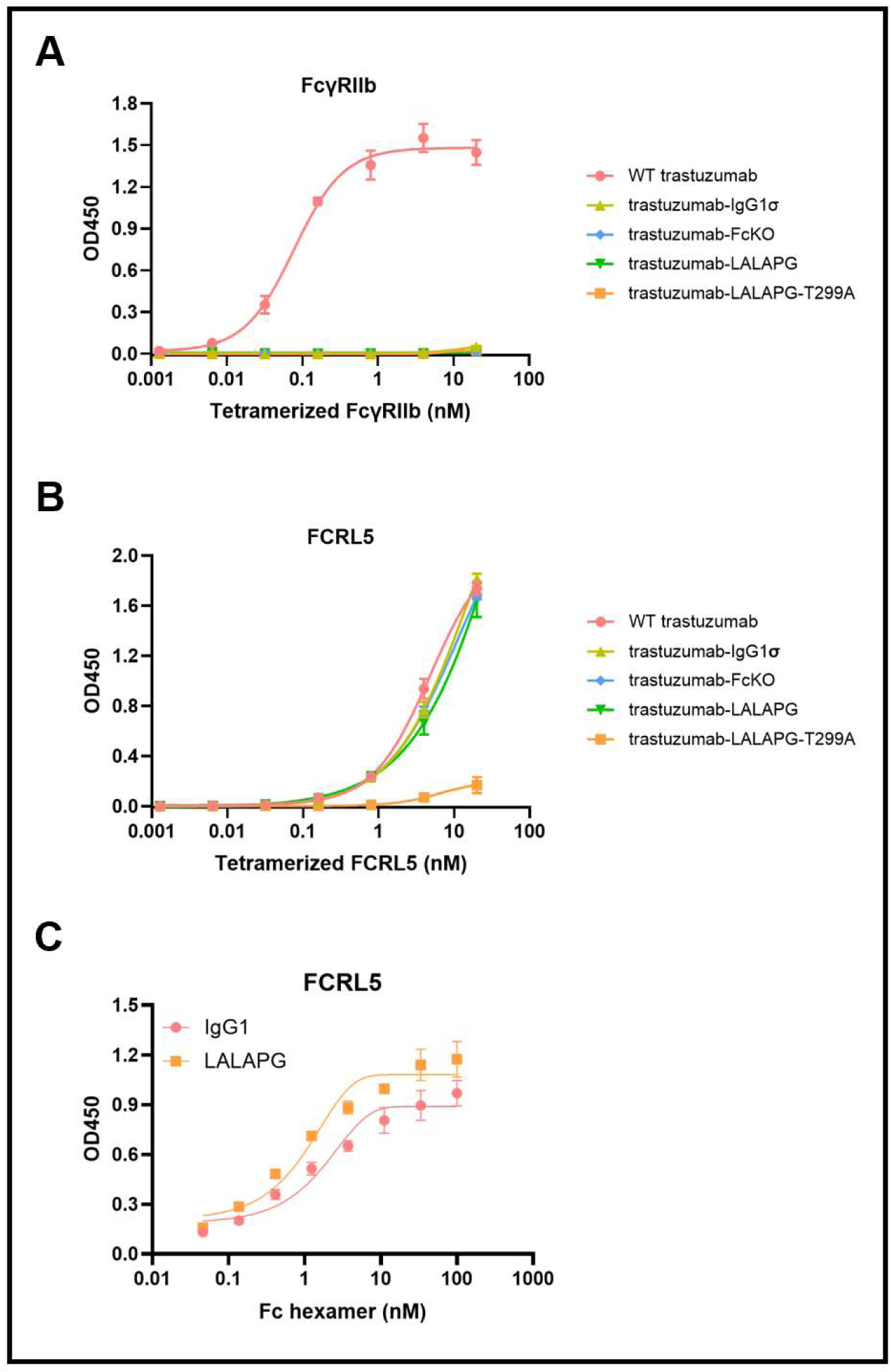
Binding of Fc-silenced IgG variants to FCRL5. Binding of Fc-silenced trastuzumab variants to (A) tetramerized FcγRIIb and (B) tetramerized FCRL5 (Q16-G851) (mean ± SD). (C) Binding of WT IgG1 Fc hexamer and IgG1 Fc-LALAPG hexamer to FCRL5 (Q16-R844) (mean ± SD).

Upon binding to surface antigen at an optimal density, IgG Fc domains can form hexamers that, in turn, engage C1q to initiate activation of the classical complement pathway (*29*). We wondered whether the proximity of the Fc domains in hexamers might affect FCRL5 binding. IgM tailpiece–fused IgG1 Fc hexamers bury substantial portions of the Fc surface during oligomerization (*30*), yet FCRL5 bound efficiently to Fc hexamers containing either wild-type (WT) IgG1 Fc or IgG1 Fc-LALAPG (Figure 1C).

### Structural analysis of the human FCRL5-IgG1 Fc complex

The inherently weak binding of FCRL5 to IgG Fc (*K*_D_ = ∼46-86 μM) poses a significant technical challenge for determining the FCRL5:Fc structure, given the very rapid dissociation of the complex. To address this limitation, we used AlphaFold-based structural modeling (*31*, *32*) to identify a plausible binding mode between FCRL5 and IgG1 Fc that is consistent with the findings presented above (Figure S2A). We then carried out alanine-scan mutagenesis on the IgG1 Fc domain to evaluate the predicted binding interface. Substitution of Fc residues D280, E293, E294, E318, E333, and Y373 with Ala reduced the binding of the FCRL5 ectodomain to trastuzumab (Figures 2A and 2B). The Y296A mutation showed no detectable binding, highlighting a critical role for this residue in FCRL5 binding. Unexpectedly, trastuzumab-Q342A bound to FCRL5 with a *K*_D_ of 6.2 ± 0.4 μM, a 2-fold improvement over WT trastuzumab. Separately, we performed a limited glutamic acid scanning mutagenesis on the Fc to enhance electrostatic interactions with FCRL5 (Figure S2B). Whereas D312E, N315E, and A330E either had no effect or slightly reduced binding, the D280E, G281E, V282E, and T335E substitutions conferred a modest increase in affinity (Figure 2C). Importantly, K317E and K320E resulted in a 7-fold and 56-fold reduced *K*_D_ (2.0 ± 0.04 µM and 0.23 ± 0.002 µM, respectively). When the K320E mutation was combined with Q342A, the effect of the two substitutions was additive, further lowering the *K*_D_ to 127 ± 1 nM (Figure 2D).

**Figure 2:**
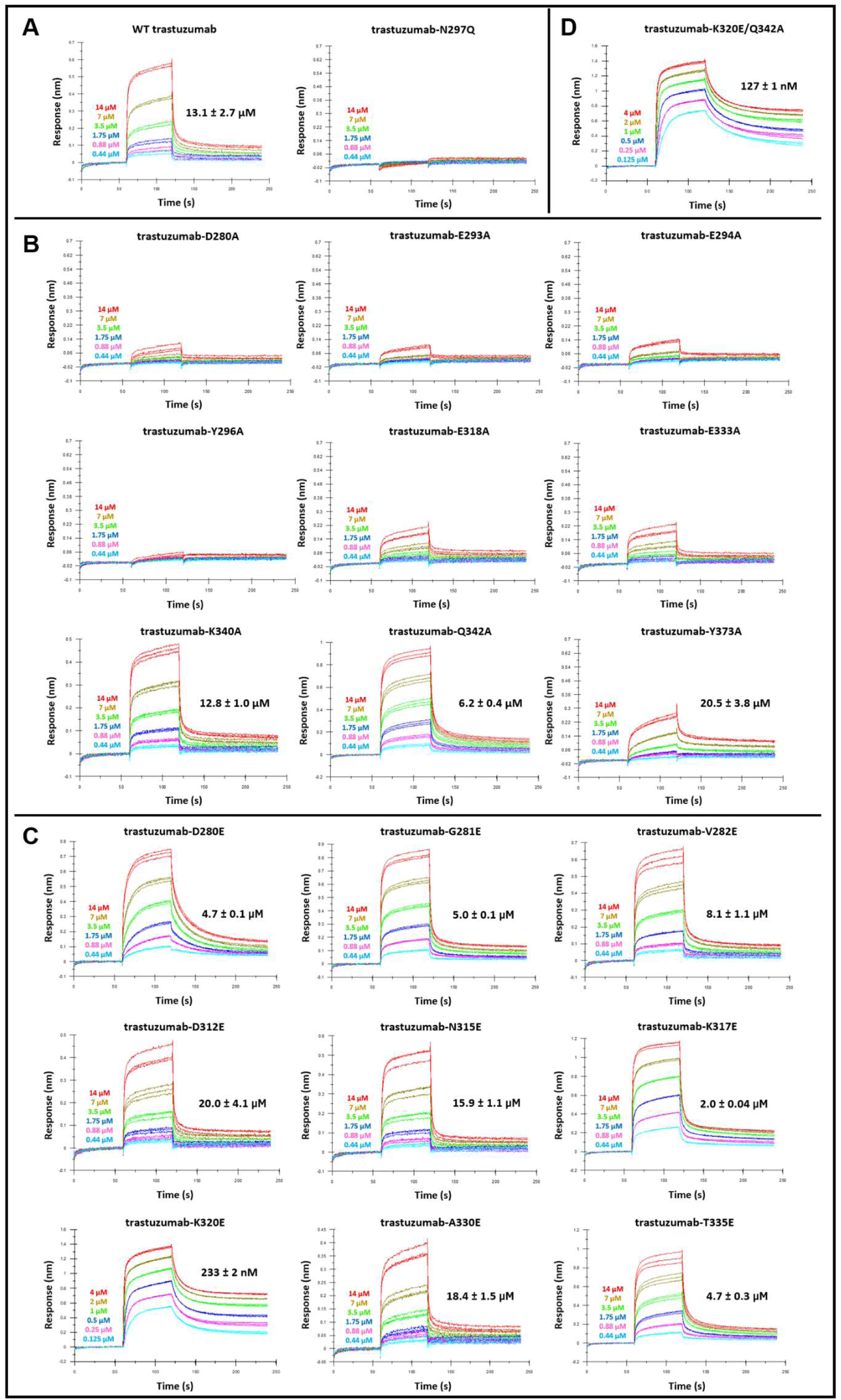
Ala- and Glu-scanning mutagenesis of the predicted IgG1 Fc interface. (A) BLI sensorgrams of WT trastuzumab and aglycosylated trastuzumab (trastuzumab-N297Q) binding to FCRL5 (Q16-G851). (B) BLI analysis of the effect of Ala substitutions on the FCRL5 affinity of trastuzumab. (C) BLI analysis of the effect of Glu substitutions. (D) BLI sensorgram of trastuzumab-K320E/Q342A binding to FCRL5 (Q16-G851). Antibodies were 2-fold serially diluted in kinetics buffer (PBS, 1% BSA, 0.02% Tween 20) starting at either 4 μM or 14 μM. Triplicate measurements were performed for each antibody, except for WT trastuzumab, for which duplicate measurements were performed. The dissociation constant (*K*_D_) is shown for each antibody as mean ± SD.

FCRL5 D1–D3 expresses very poorly in Expi293F cells (Figure S3D), analogous to the very low expression of the FcγRI ectodomain, which similarly comprises three Ig-like domains (*33*). Inspired by the work of Ide and coworkers, who used directed evolution to produce a well-expressed variant of the FcγRI ectodomain (*33*) that, in turn, enabled the production of a crystallizable complex for structural studies (*34*), we used directed evolution to isolate an FCRL5 D1-D3 variant with enhanced expression in Expi293F cells (Figure S3). This was achieved by mutagenizing the gene encoding the first three extracellular domains of FCRL5 using error-prone PCR, followed by display on *S. cerevisiae* (Figure S3A). After four rounds of enrichment, we evaluated the surface expression of individual FCRL5 D1-D3 clones on yeast (Figures S3B and S3C), selected the five best-performing clones, and assessed their recombinant expression in Expi293F cells (Figure S3D). Two clones that showed markedly higher expression levels in Expi293F cells compared to WT FCRL5 D1–D3 were obtained, and the mutations were combined to create rFCRL5 D1-D3 (Figure S3E). rFCRL5 D1-D3 contains a total of six amino acid substitutions (F26S, K45E, L111P, E128G, H155R, and W251R), was expressed at approximately 0.03 mg/mL in Expi293F cells, and binds to WT trastuzumab and the trastuzumab-K320E/Q342A variant with the expected affinities (Figures S3F–S3H).

rFCRL5 D1-D3 and IgG1 Fc-K320E/Q342A were each purified using affinity chromatography followed by size-exclusion chromatography (SEC) on a Superdex 200 column (Figures S3F and S3G, S4A and S4B). When rFCRL5 and IgG1-Fc K320E/Q342A were mixed in a 1:2 molar ratio, the reconstituted sample eluted as two partially overlapping peaks (Figure S5C) with the first peak having an apparent molecular weight (M.W.) of 83 kDa (Figure S5D), consistent with a 1:1 FCRL5-Fc complex. The second peak corresponded to an apparent M.W. of 48 kDa, as expected for excess unbound Fc. Notably, no higher-molecular-weight species were observed despite the presence of excess Fc, consistent with a 1:1 binding stoichiometry.

To further examine the stoichiometry of the FCRL5-IgG complex, we performed native mass spectrometry at varying protein ratios. Under native-like conditions, trastuzumab- K320E/Q342A exhibited a well-resolved charge state distribution spanning the 19+ to 25+ charge states, yielding a deconvoluted mass of 149.9 kDa consistent with its expected M.W. of 149.5 kDa (Figure S5E). rFCRL5 D1-D3 was detected predominantly in the 9+ to 13+ charge states, resulting in a deconvoluted mass of 33.6 kDa (Figure S5F). When rFCRL5 D1-D3 and trastuzumab-K320E/Q342A were mixed at equimolar concentrations (5 μM each), a new set of higher-mass species was evident in the m/z range of ∼6000–7000 (Figure S5G). Deconvolution of these ion peaks yielded a mass of 183.3 kDa, consistent with formation of a 1:1 FCRL5-IgG complex. Mixing rFCRL5 D1-D3 and trastuzumab-K320E/Q342A at a 10:1 molar ratio (9.1 μM and 0.9 μM, respectively) also yielded a 1:1 complex, with no evidence of 2:1 or higher-order complexes between the two proteins (Figure S5H).

We crystallized the rFCRL5 D1-D3:IgG1 Fc-K320E/Q342A complex and solved the structure. Diffracting crystals were indexed to a resolution of 3.4 Å in space group P3₁21 with a single complex molecule per asymmetric unit (Figure 3A; Table S1). Continuous electron density was observed for FCRL5 residues R22–I282, with the exception of residues F47–P52 in domain 1, which were not resolved due to conformational flexibility. For IgG1 Fc, well-defined electron density extended from G237 to P445 in chain A and from L235 to P445 in chain B. In the refined structure, both Fc chains contain a core-fucosylated, agalactosylated biantennary complex N-glycan (G0F) at N297 (Figure 3B). However, one terminal N-acetylglucosamine (GlcNAc) was not resolved for the glycan attached to chain A. FCRL5 is glycosylated at N132. Clear density was observed for the first GlcNAc residue (Figure 3B).

**Figure 3.**
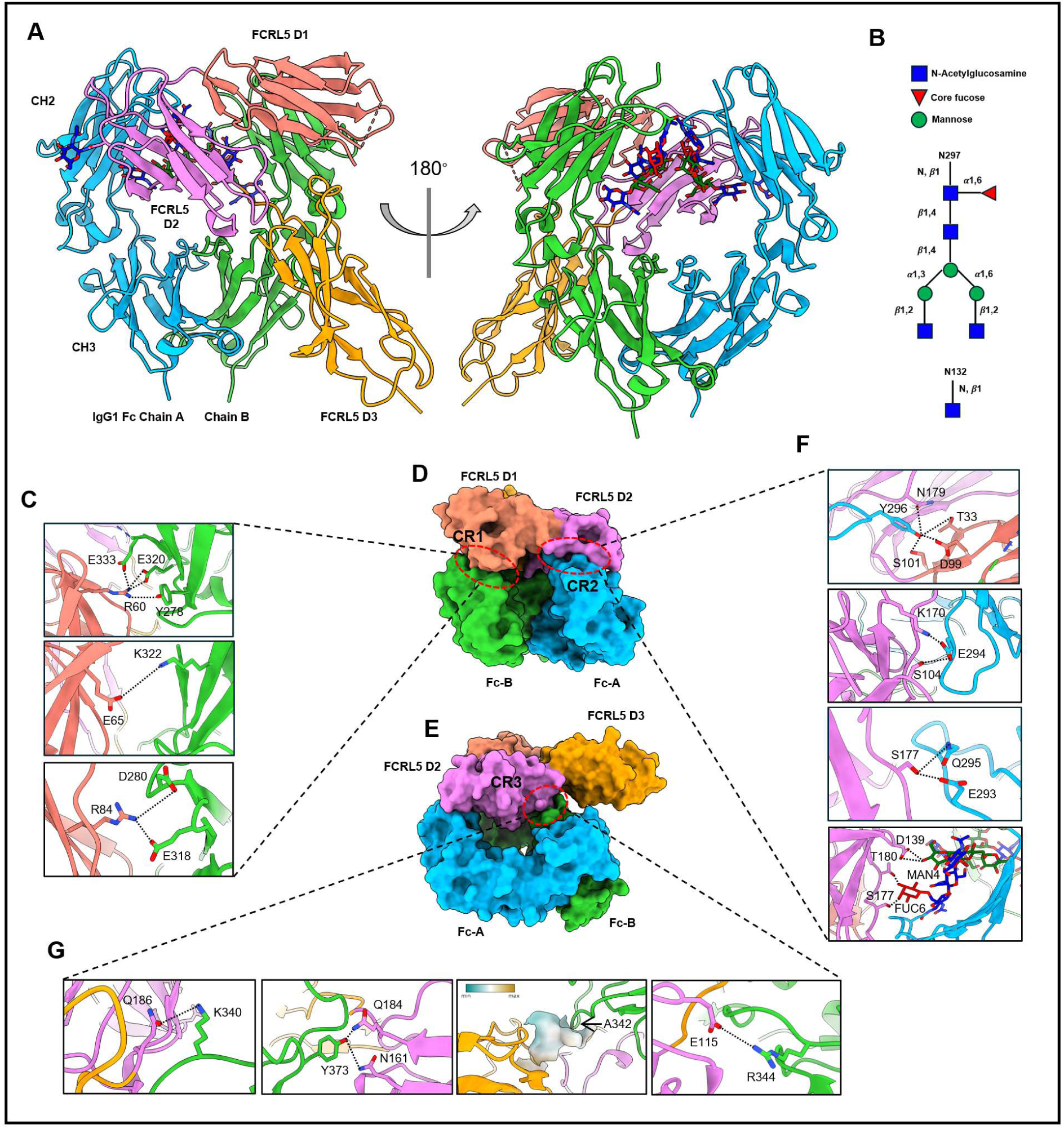
Structural characterization of the FCRL5–IgG1 Fc complex and interaction interfaces. (A) Crystal structure of rFCRL5 D1-D3 in complex with IgG1 Fc-K320E/Q342A shown in ribbon representation. IgG1 Fc chains A and B are colored light blue and light green, respectively, while FCRL5 domains D1, D2, and D3 are shown in salmon, violet, and orange. The CH2 and CH3 domains of Fc are indicated. A 180° rotated view illustrates the overall architecture of the complex and the mode of Fc engagement by FCRL5. (B) Schematic representation of N-linked glycans on IgG1 Fc (N297) and FCRL5 (N132). (C) Close-up views of representative electrostatic and hydrogen-bond interactions at contact region 1 (CR1) between FCRL5 D1 and Fc. Key residues contributing to interface stabilization are shown as sticks. (D) Surface representation of the complex illustrating interaction interfaces between FCRL5 and Fc. Two contact regions (CR1 & CR2) are highlighted, showing engagement of FCRL5 D1-D2 with both Fc chains. (E) Alternate orientation emphasizing contact region 3 (CR3), which is formed by interactions between the FCRL5 D2/D3 interface and the CH3 domain of Fc chain B. (F) Detailed views of contact region 2 (CR2) showing residue-specific interactions and glycan-proximal contacts contributing to receptor recognition. (G) Additional interface close-ups of CR3. Dashed lines indicate locations of magnified regions shown in panels 3C, 3F, and 3G.

FCRL5 adopts a canonical immunoglobulin fold comprising three C2-type Ig-like domains (Figure 3A). Of note, structural analysis confirmed that none of the amino acid substitutions that enhance the expression of rFCRL5 D1-D3 are located at the binding interface with IgG1 Fc (Figure S6). FCRL5 D1-D3 interact with the Fc through three distinct contact regions (CRs) in a front-facing configuration (Figures 3A, 3D, and 3E). The D1 domain of FCRL5 forms multiple salt bridges with the CH2 domain of Fc chain B, defining contact region 1 (CR1; Figure 3C). FCRL5 domains D1 and D2 engage residues of the C’E loop (E293-Y296) and the N297-linked glycan on Fc chain A, forming contact region 2 (CR2; Figure 3F). The third contact region (CR3) is formed by interactions between the FCRL5 D2/D3 interface and the CH3 domain of Fc chain B (Figure 3G). FCRL5 thus binds the Fc asymmetrically, with approximately 654 Å² of surface area buried via interactions with Fc chain B and roughly 380 Å² via Fc chain A. Within CR1, the K320E substitution in IgG1 Fc converts local electrostatic repulsion or unsatisfied charge into a stabilizing network of salt-bridge and hydrogen-bonding interactions that anchor the Fc to FCRL5 (Figure 3C). Fc residues D280 and E318 form salt bridges with FCRL5 residue R84. FCRL5 residue E65 further contributes to CR1 stabilization through a salt bridge with Fc residue K322 (Figure 3C). CR2 is stabilized by an extensive interaction network comprising both protein–protein and glycan–protein contacts (Figure 3F). Y296 from Fc chain A engages residues from FCRL5 D1 and D2, including N179,

T33, D99, and S101, forming multiple hydrogen bonds (Figure 3F). The FCRL5 residue K170 forms a salt bridge with Fc residue E294, which also engages in an additional hydrogen bond with FCRL5 residue S104. Fc residues E293 and Q295 form hydrogen bonds with S177 of FCRL5. In addition, the biantennary complex N-glycan at N297 of Fc chain A interacts with FCRL5 domain D2, where the core fucose (Fuc6) and the mannose residue from one antenna (Man4) form hydrogen bonds with D139, T180, and S177 of FCRL5 D2, further strengthening the interaction (Figure 3F). The glycan:D2 interaction is likely part of the reason why the N297Q substitution markedly reduces FCRL5 binding (Figure 2A), with destabilization of the C’E loop likely also contributing (*35*). The third contact region involves hydrogen bonding, electrostatic interactions, and hydrophobic interactions, which together help stabilize the FCRL5–Fc interface (Figure 3G). FCRL5 residue Q186 creates a hydrogen bond with K340 of Fc chain B. The Fc residue Y373 forms hydrogen bonds with N161 and Q184 of FCRL5. The Q342A substitution in IgG1 Fc removes a polar glutamine and extends a contiguous hydrophobic motif (G341–A342–P343) that packs against L158 and V185 of FCRL5, thereby improving van der Waals complementarity and eliminating an energetically unfavorable polar group at the interface (Figure 3G). FCRL5 residue E115 forms a salt bridge with R344 of Fc chain B. Collectively, the cooperative contributions of CR1–CR3 establish a unique Fc recognition mode for FCRL5 that is clearly distinct from that of any other Fc-binding protein (Figure 4A).

**Figure 4.**
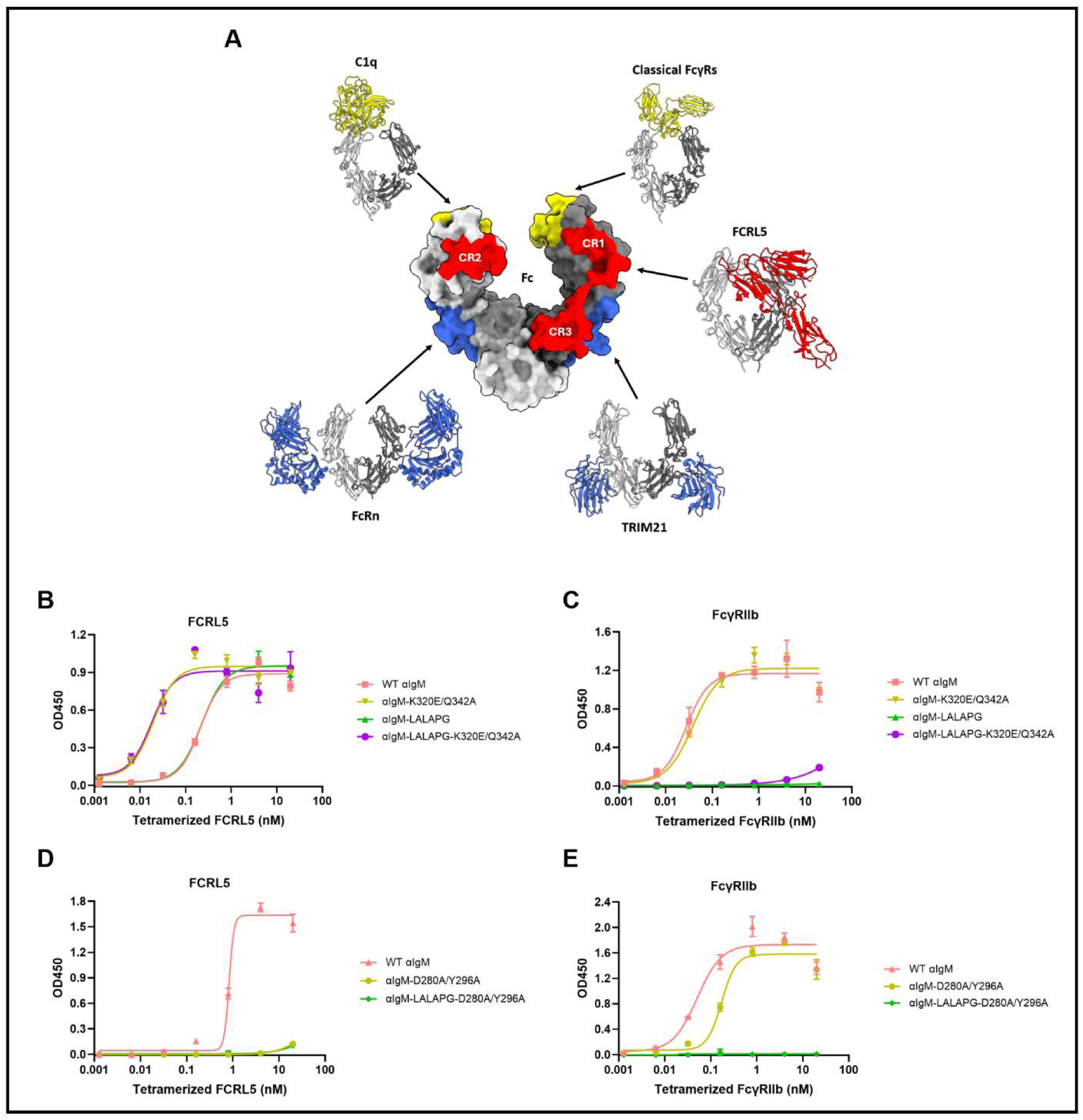
The FCRL5 affinity of IgG1 can be selectively tuned with minimal impact on FcγRIIb affinity. (A) FCRL5 binds to a site on the IgG1 Fc domain that is distinct from the binding site of classical FcγRs (PDB: 4W4O), C1q (PDB: 6FCZ), FcRn (PDB: 7Q15), or TRIM21 (PDB: 2IWG). The surface of the Fc domain in the center shows the contact area of each Fc-binding protein. The Fc-binding proteins and their contact areas are colored based on where they bind the IgG Fc domain. C1q and the classical FcγRs bind IgG Fc at the lower hinge and upper CH2 domain (yellow), while FcRn and TRIM21 engage IgG Fc at the CH2- CH3 interface (blue). FCRL5 interacts with the IgG Fc domain through three distinct contact regions (red). CR1 = contact region 1, CR2 = contact region 2, and CR3 = contact region 3. (B-E) ELISA measurements of Fc-engineered anti-human IgM (αIgM) antibody variants binding to (B, D) tetramerized FCRL5 (Q16-G851) and (C, E) tetramerized FcγRIIb (mean ± SD).

### Dissecting the effect of FCRL5 and FcγRIIb on B cell calcium flux upon BCR co-ligation

Unlike classical FcγRs and C1q, which bind IgG Fc at the lower hinge and upper CH2 domain, and unlike FcRn and TRIM21, which target the CH2–CH3 interface, FCRL5 engages an extensive binding interface on the viewer-facing side of IgG Fc (Figure 4A). Apart from FCRL5, FcγRIIb is the only other Fcγ receptor found on B cells (*2*). Agonism of FcγRIIb by antibodies that also engage the BCR complex is well established to inhibit Ca^2+^ flux, reduce proliferation and differentiation, and has been exploited in clinical-stage experimental therapeutics for certain autoimmune indications (*36*–*39*). Although previous studies have shown that co-ligation of FCRL5 with the BCR inhibits Ca^2+^ flux (*19*–*21*), this has not yet been demonstrated under conditions in which the Fc region of IgG is used to agonize WT FCRL5, which would more accurately reflect physiological conditions. Therefore, we utilized the data from our structural analyses and biochemical studies outlined above to develop a panel of Fc-engineered monoclonal anti-human IgM (αIgM) antibodies. These IgG1 antibodies can cross-link IgM BCRs through their Fab regions and selectively engage FCRL5, FcγRIIb, or both, depending on their engineered Fc region. As expected, WT αIgM bound to both FCRL5 and FcγRIIb, while the LALAPG substitutions in αIgM-LALAPG ablated only FcγRIIb binding (Figures 4B and 4C). Introduction of the K320E/Q342A substitutions into WT αIgM and αIgM- LALAPG markedly lowered their EC50 for FCRL5, while their EC50 for FcγRIIb was essentially unaffected (Figures 4B and 4C). Introduction of D280A/Y296A substitutions into WT αIgM and αIgM-LALAPG nearly abolished binding to FCRL5 while minimally affecting binding to FcγRIIb (Figures 4D and 4E).

We used the antibody panel above to assess how co-ligating either or both Fc receptors with the BCR affects Ca^2+^ flux in the well-studied diffuse large B cell lymphoma cell line, TMD8 (*40*). We first quantitatively determined the relative expression of FCRL5 and FcγRIIb on TMD8 cells. In brief, TMD8 cells and bead populations with known antibody binding capacity (ABC) were stained with fluorescent anti-FCRL5 or anti-FcγRIIb mAbs. Isotype controls verified that the staining antibodies did not bind to the Fc receptors via their Fc regions. ABC values provide an approximate estimate of the number of Fc receptors present on TMD8 cells, as each staining antibody can bind up to two Fc receptors due to its bivalency. The surface expression of FcγRIIb on TMD8 cells is approximately 9x higher than that of FCRL5, with ABC values of ∼33,400 ± 700 and ∼3,700 ± 300, respectively (Figure 5A). As expected, cross-linking the BCR with αIgM-LALAPG-D280A/Y296A, which cannot engage either FCRL5 or FcγRIIb, resulted in more sustained Ca^2+^ flux than cross-linking the BCR with WT αIgM, which binds to both Fc receptors (Figure 5B). BCR ligation using the αIgM-D280A/Y296A antibody, which binds only to FcγRIIb, resulted in significant inhibition of Ca^2+^ flux relative to the negative control αIgM-LALAPG D280A/Y296A, as expected (Figure 5C). Selective co-ligation of the BCR with FCRL5 by αIgM-LALAPG also resulted in a small but reproducible reduction of Ca^2+^ flux relative to αIgM-LALAPG D280A/Y296A (Figure 5D). The smaller effect of FCRL5 engagement relative to that of FcγRIIb may, in part, be due to the much lower expression of FCRL5 on TMD8 cells (Figure 5A). The higher-affinity variant, αIgM-LALAPG- K320E/Q342A, showed much stronger inhibition of Ca^2+^ flux (Figure 5D). Enhancing FCRL5 affinity in WT αIgM also resulted in stronger inhibition of Ca^2+^ flux (Figure 5E), highlighting that even in the presence of strong FcγRIIb signaling, increased FCRL5 agonism further inhibits Ca^2+^ flux in B cells. Taken together, these data demonstrate that Fc-mediated recruitment of FCRL5 to cross-linked BCRs inhibits Ca^2+^ flux in FCRL5-expressing B cells. This effect is independent of and distinct from that of FcγRIIb engagement, and the inhibition of Ca^2+^ flux is enhanced by mutations that increase FCRL5 affinity.

**Figure 5.**
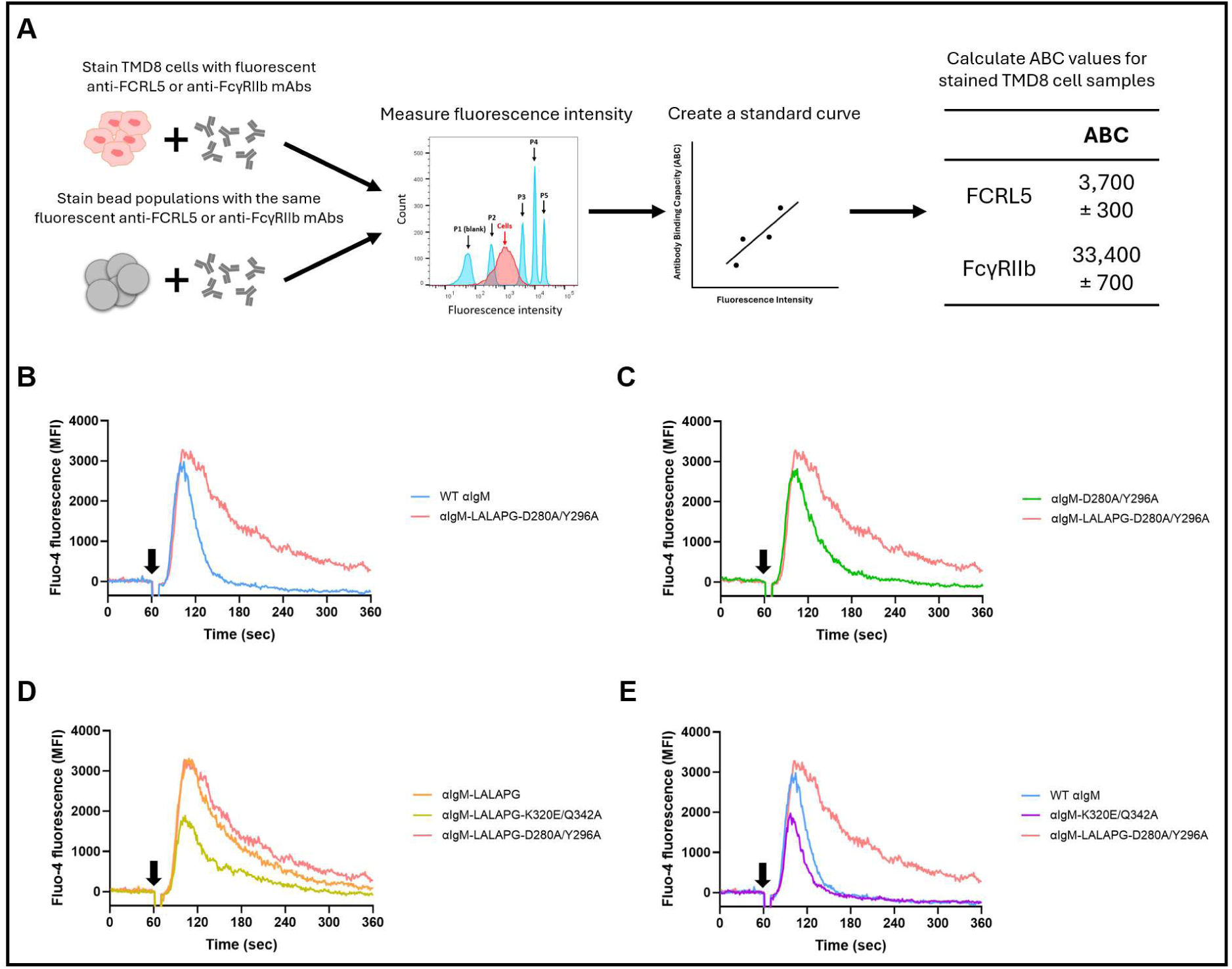
Fc-mediated recruitment of FCRL5 to cross-linked BCRs inhibits BCR-mediated calcium mobilization *in vitro*. (A) Quantification of the surface expression of FCRL5 and FcγRIIb on TMD8 cells. Antibody binding capacity (ABC) values are presented as mean ± SD. (B-E) Flow cytometric analysis of the intracellular calcium levels in TMD8 cells stimulated with Fc-engineered αIgM antibodies. TMD8 cells were loaded with the calcium indicator Fluo-4 AM. Baseline was measured for 60 seconds. Fc-engineered αIgM antibodies were added to the TMD8 cells at t = 60 sec (black arrow). Calcium mobilization was measured for five minutes.

## Discussion

We observed that IgG1 engineered with mutations that silence Fc effector functions, such as LALAPG, IgG1σ, and FcKO, which are widely used in clinical-stage and approved antibody therapeutics, retain binding to the FCRL5 receptor on B cells. The structure and function of FCRL5 in B cell biology are largely unexplored and remain enigmatic. Earlier studies have suggested that FCRL5 can have both inhibitory and activating functions (*19*–*21*), and its expression varies greatly during B cell development (*41*). Despite the paucity of clear information on its biological role, FCRL5 has emerged in recent years as a therapeutic target of major significance. Most importantly, in multiple myeloma (MM) FCRL5 is expressed at high levels with near 100% penetrance (*23*, *24*), and a recent study suggests that it is also a key surface marker for MM-initiating cells (*25*). These observations have provided the rationale for developing FCRL5-targeting therapeutics for MM, including antibody-drug conjugates, anti-FCRL5 CAR T cells, and bispecific antibodies that co-engage CD3, which are currently being evaluated in multiple clinical studies or are in advanced preclinical development, with cevostamab currently in mid-stage phase III trials (*25*, *42–44*).

Beyond MM, FCRL5 may be clinically relevant in multiple other disease settings. FCRL5 expression has been reported on subsets of chronic lymphocytic leukemia, mantle cell lymphoma, and hairy cell leukemia patients (*18*, *45*, *46*). Separately, in myasthenia gravis, FCRL5 expression occurs across all B cell subsets, with the highest level observed on age-associated B cells, which become expanded in relapsed patients following rituximab or BCMA/CD19 CAR T cell therapy (*47*). CD11c^hi^ B cells are expanded in SLE and express high levels of FCRL5 (*48*). Single-nucleotide polymorphisms in FCRL5 are linked to multiple sclerosis (MS) susceptibility (*49*), and recent evidence indicates that soluble FCRL5 levels in the cerebrospinal fluid of MS patients may serve as a biomarker for diagnosis and disease activity prediction (*50*). Epstein-Barr virus infection is the primary environmental risk factor for MS (*51*), and it is established that in infected B cells, EBNA2 induces FCRL5 expression in a CBF1-dependent manner (*52*). High FCRL5 expression is also a hallmark of atypical memory B cells, which exhibit an anergic phenotype and are resistant to BCR-mediated activation (*53*, *54*). The consequences of FCRL5 engagement by immune complexes formed by Fc-silenced therapeutic antibodies in these disease populations, and more broadly on FCRL5+ long-lived bone marrow cells, are unknown. In addition to a direct effect on B cell signaling, immune complexes bound to FCRL5 are internalized by B cells and processed in lysosomes (*15*, *24*). FCRL5-mediated internalization of immune complexes may play a role in antigen presentation and, possibly, in the immunogenicity of therapeutic antibodies (*55*). Overall, our findings highlight an unappreciated biological effect of Fc-silent antibodies on B cell function that warrants further investigation.

The low affinity of the IgG1 Fc domain for FCRL5, coupled with the poor expression of the ligand-binding portion of the receptor, the D1-D3 domains, has hindered the structural analysis of the FCRL5:Fc interaction. Here, we addressed these two limitations by: (i) using directed evolution and yeast display to isolate rFCRL5 D1-D3, which could be expressed at preparative levels in Expi293F cells, and (ii) structural modeling coupled with Ala and Glu mutagenesis to engineer IgG1 Fc-K320E/Q342A, which has ∼100-fold increased affinity for FCRL5. In this manner, we succeeded in solving the crystal structure of FCRL5 D1-D3 in complex with IgG1 Fc at 3.4 Å resolution. We show that FCRL5 binds to the IgG1 Fc domain at a unique site, distinct from the binding sites of other Fc-binding proteins such as the classical FcγRs, C1q, FcRn, and TRIM21 (Figure 4A). Structurally, FCRL5 recognition is distinguished by its asymmetric engagement of both Fc chains through three spatially separated contact regions, integrating electrostatic, hydrogen-bonding, and glycan-mediated interactions. Notably, the interface extensively involves residues distal from the canonical FcγR and C1q binding sites, explaining why Fc-silencing mutations such as LALAPG do not disrupt FCRL5 binding.

While this manuscript was in preparation, Xiao and coworkers reported that a stabilized IgG1 Fc hexamer, stabilized by fusing a modified IgM tailpiece to the IgG1 Fc and by introducing an L309C mutation that participates in a disulfide bond with a proximal Fc, binds to FCRL5 with an apparent *K*_D_ in the 200 nM range (*15*). They then used single-particle cryo-EM analysis to solve the structure of the IgG1 Fc hexamer in complex with FCRL5. Xiao *et al.* reported that FCRL5 D1-D3 bind to IgG Fc in a 1:2 ratio stoichiometry. Superposition of our crystal structure with the cryo-EM structure of Xiao *et al.* (PDB: 9LOC) showed minimal overall structural differences with a Cα RMSD of 0.84 Å for IgG1 Fc and 0.90 Å for FCRL5 (Figure S7A). However, Xiao *et al.* reported a second interface between FCRL5 domain D3 and the CH3 domains of a neighboring Fc in the planar hexamer. At the reported interface between FCRL5 D3 and the second Fc molecule in the cryo-EM structure, the proposed hydrogen bonds between FCRL5 residue Q200 and Fc residues N389 and S400, as well as the reported van der Waals interactions between Fc residue R416 and FCRL5 residues F257 and W277, are geometrically compatible with our crystal structure (Figures S7B and S7C). However, our crystal structure does not contain electron density corresponding to a second Fc molecule, despite the presence of these potentially permissive interaction surfaces. Moreover, native mass spectrometry and SEC also support a 1:1 stoichiometry in solution, indicating that the 1:2 ratio observed in the cryo-EM structure likely reflects avidity-driven interactions within a high-density Fc environment rather than the intrinsic binding mode. Also, the fact that heat-aggregated IgG, which is not ordered, binds to FCRL5 suggests that high-avidity interactions can drive FCRL5 binding without the requirement for a strict planar arrangement as that found in a hexamer (*12*, *15*). Conversely, however, we acknowledge that our work involves engineered high-affinity Fc domains and rFCRL5 D1-D3, which may play a role in dictating the 1:1 stoichiometry we observed.

These details aside, both the crystal structure presented here and the cryo-EM structure by Xiao *et al.* demonstrate that FCRL5 binds to IgG in a manner clearly distinct from that of all other known Fc-binding proteins. We then demonstrated that Fc-silenced IgG can be used to selectively co-ligate FCRL5 with the BCR *in cis*, resulting in the inhibition of Ca^2+^ flux in FCRL5-expressing B cells. This inhibition is independent of and distinct from the inhibition caused by FcγRIIb engagement. Enhanced FCRL5 affinity resulted in stronger inhibition of BCR-induced Ca^2+^ flux, even in the presence of FcγRIIb signaling. IgG antibodies that selectively engage FCRL5 with varying affinity could be highly useful for future mechanistic research, as they could aid in understanding the physiological role of FCRL5 on atypical memory B cells and plasma cells in both health and disease.

## Materials and methods

### Cells and Reagents

TMD8 cells were obtained from Cells Online LLC. Cells were cultured in RPMI 1640 with 10% fetal bovine serum (FBS) and 1x Penicillin-Streptomycin. Cells were passaged every 2-3 days at a seeding density of 0.1 - 0.2 x 10^6 cells/mL. Recombinant human FCRL5 protein was purchased from RCD Systems (cat. no. 2078-FC). Biotinylated human FCRL5 protein was purchased from ACROBiosystems (cat. no. FC5-H82E3) and Creative BioMart (cat. no. FCRL5-971HB). Biotinylated human FcγRIIb protein was purchased from ACROBiosystems (cat. no CDB-H82E0). Bovine Serum Albumin (BSA) was purchased from Sigma-Aldrich (cat. no. A3294). Ammonium bicarbonate and LC-MS-grade water were purchased from Sigma-Aldrich (St. Louis, MO). Micro Bio-Spin P6 gel columns were obtained from Bio-Rad Laboratories (Hercules, CA, USA).

### Preparation of antibodies and Fc hexamers

pcDNA3.4-trastuzumab, a mammalian expression plasmid harboring the trastuzumab heavy chain sequence, served as the template for cloning. Site-directed mutagenesis was performed by Gibson Assembly (*56*). The monoclonal anti-human IgM (αIgM) antibodies were made by substituting trastuzumab’s variable regions with the variable regions of a mouse anti-human IgM mAb (*57*). To generate the Fc hexamers, a L309C mutation was added to the Fc domain of IgG1, and the IgM μ-tailpiece (PTLYNVSLVMSDTAGTCY) was fused to the C-terminus of the Fc domain (*58*, *59*). V567I and A572G were introduced into the IgG1 Fc- LALAPG hexamer to increase the stability of the hexameric structure and enhance the expression yield (*30*).

Antibodies and Fc hexamers were produced by transient transfection of Expi293F cells (Thermo Fisher Scientific, cat. no. A14527). Cells were cultured on an orbital shaker platform (120 rpm) inside a temperature and CO2-controlled incubator (37 °C, 8% CO_2_). Antibodies and Fc hexamers were harvested five days after transfection. Antibodies and Fc hexamers were purified by affinity chromatography using protein G agarose resin (Thermo Fisher Scientific, cat. no. 20399). 100 mM glycine-HCl (pH 2.7) was used for elution. The eluate was immediately neutralized with 1 M Tris-HCl (pH 8.0), and the samples were buffer-exchanged into PBS. The purity and oligomeric state of all antibodies and Fc hexamers were assessed by SDS-PAGE and size-exclusion chromatography.

### Engineering of rFCRL5 D1-D3

The yeast display library was constructed according to the protocol described in (*60*). *Saccharomyces cerevisiae* strain AWY101 was used as the host strain (*61*). Inserts were generated by error-prone PCR (*62*) of wild-type FCRL5 cDNA at a 0.2% error rate and introduced together with BamHI/NdeI-linearized pCTCON2 by electroporation. The library size (2 x 10^7^ mutants) was determined by colony counting. Selected colonies were sequenced to assess library diversity. Protein expression was induced in phosphate-buffered SG-CAA medium (20 g/L galactose, 6.7 g/L YNB without amino acids, 5 g/L Casamino Acids, 5.4 g/L Na_2_HPO_4_, and 8.6 g/L NaH_2_PO_4_ · H_2_O). Induced cells were labeled with anti-FCRL5-PE antibody (BioLegend, cat. no. 340304) in PBS containing 0.5% BSA and 2 mM EDTA, then sorted using a BD FACSAria IIu cell sorter. In total, the library underwent four rounds of sorting. After the fourth round, 17 mutant clones and 3 WT controls were analyzed individually for FCRL5 expression. Each clone was assigned an expression score calculated as the percentage of cells in the high-expression quadrant (Q3) multiplied by the median fluorescence intensity (MFI) of that quadrant. Based on these scores, the five best-performing mutants were subcloned into a mammalian expression vector (pcDNA3.4) and evaluated for expression in Expi293F cells. The two mutants that showed markedly higher expression levels in Expi293F cells compared to WT FCRL5 D1–D3 were combined to create rFCRL5 D1-D3.

### Preparation of rFCRL5 D1-D3 and IgG1 Fc-K320E/Q342A

IgG1 Fc-K320E/Q342A (EU numbering: D221-K447) and rFCRL5 D1-D3 (Q16-I282) were each cloned into pcDNA3.4 vectors by Gibson assembly, and proteins were expressed by transient transfection of Expi293F cells as described above. IgG1 Fc-K320E/Q342A was isolated by Protein G chromatography. rFCRL5 D1-D3 was isolated using a Ni-NTA column. Supernatants were loaded three times to ensure that all protein was bound to the column. Protein G columns were washed twice with PBS, and IgG1 Fc-K320E/Q342A was eluted using 100 mM glycine-HCl (pH 2.7). The eluate was immediately neutralized by the addition of 1 M Tris-HCl (pH 8.0). For Ni-NTA purification, the resin was washed sequentially with 10 mM imidazole buffer (pH 7.4) and then with 20 mM imidazole buffer (pH 7.4). rFCRL5 D1-D3 was eluted using a 300 mM imidazole buffer (pH 7.4). Both proteins were buffer-exchanged into PBS and concentrated using Vivaspin® centrifugal concentrators. The purity and oligomeric state of rFCRL5 D1-D3 and IgG1 Fc-K320E/Q342A were assessed by SDS-PAGE and size-exclusion chromatography.

### Size exclusion chromatography

SEC analysis of all proteins was performed on an ÄKTA pure system (GE healthcare) using a Superdex 200 Increase 10/300 GL column (Cytiva, cat. no. 28990944). The mobile phase was PBS at a flow rate of 0.75 mL/min.

### Structure predictions

Initial protein structure predictions of the complex between FCRL5 and IgG1 Fc were made using ColabFold (*31*). The query sequence would consist of one or two copies of the first three N-terminal domains (Q16-I282) of human FCRL5 (UniProt accession number: Q96RD9-1) and two copies of a portion (D104-K330) of human immunoglobulin heavy constant gamma 1 (IGHG1; UniProt accession number: P01857-1). Later protein complex predictions were made using the AlphaFold 3 server (*32*). In addition to the query sequence described above, AlphaFold 3 predictions were informed of the presence of a glycan (N-Acetyl-beta-D-glucosamine) attached to N180 (EU numbering: N297) of IGHG1.

### Biolayer interferometry

BLI assays were performed on an Octet RED96e System (FortéBio). Biotinylated human FCRL5 protein (ACROBiosystems, cat. no. FC5-H82E3) was loaded onto Octet High Precision Streptavidin (SAX) Biosensors (Sartorius, cat. no. 18-5117) until a threshold of 2.5 nm. Antibodies were 2-fold serially diluted in kinetics buffer (PBS, 1% BSA, 0.02% Tween 20) starting either at 4 μM or 14 μM. Triplicate measurements were performed for each antibody unless otherwise specified. Association was measured for 60 seconds, followed by dissociation for 120 seconds. Assays were performed at 25 °C while shaking at 1000 rpm. 1 M MgCl_2_ was used as the regeneration buffer. 96-well BLI plates were purchased from Greiner Bio-One (cat. no. 655209). Dissociation constants were determined using steady state analysis.

### ELISA assays

96-well plates (Corning, cat. no. 3361) were coated with antibody in PBS (10 µg/mL, 50 µL/well) and incubated overnight at 4 °C. The next morning, plates were washed three times with PBST (0.05% Tween 20 in PBS). Plates were blocked by adding 300 µL/well of 3% PBSA (3% BSA in PBST) and incubated for 2 hours at room temperature with gentle shaking. Plates were washed three times with PBST. Tetramerized Fc receptor complexes were made by mixing biotinylated Fc receptor (FCRL5: ACROBiosystems, cat. no. FC5-H82E3, FcγRIIb: ACROBiosystems, cat. no CDB-H82E0) with purified streptavidin (BioLegend, cat. no. 280302) at a 4:1 molar ratio. Tetramerized Fc receptor complexes were 5-fold serially diluted in 3% PBSA starting at 20 nM. Tetramerized complexes were added to each plate (50 µL/well) and incubated for 1 hour at room temperature with gentle shaking. Plates were washed three times with PBST. HRP anti-His tag antibody (BioLegend, cat. no. 652503) was diluted 5000x in 3% PBSA prior to use. Plates were incubated with HRP anti-His tag antibody (50 µL/well) for one hour at room temperature with gentle shaking. Plates were washed three times with PBST. 50 µL of TMB substrate (Thermo Scientific, cat. no. 34028) was added to each well, and the reaction was stopped by adding 50 µL of 2 M H_2_SO_4_ per well. Absorbance was measured at 450 nm.

Binding to IgG Fc hexamers was evaluated using FCRL5 (RCD Systems, cat. no. 2078-FC) coated at 5 μg/mL onto a high-binding 96-well plate (Corning, cat. no. 3361) and incubated overnight at 4°C. The plate was blocked with 3% BSA in PBS, and serially diluted Fc hexamers were incubated for 2 hours at RT. After washing with PBS, the plate was incubated with a 2000-fold dilution of goat anti-human IgG Fc secondary antibody, HRP (Invitrogen, cat. no. A18817). One hour later, the plate was developed with TMB substrate (Thermo Scientific, cat. no. 34028) and quenched with 2 M H_2_SO_4_. Absorbance was measured at 450 nm.

rFCRL5 D1-D3 was coated at 10 μg/mL onto a high-binding 96-well plate (Corning, cat. no. 3361) and incubated overnight at 4°C. The plate was washed three times with PBST (0.05% Tween 20 in PBS). The plate was blocked with 3% PBSA (3% BSA in PBST) and incubated for 2 hours at room temperature with gentle shaking. The plate was washed three times with PBST. Trastuzumab antibodies were added to the plate (3-fold serially diluted in 3% PBSA, starting at 5000 nM) and incubated for 1 hour at room temperature with gentle shaking. The plate was washed three times with PBST, and a 2000-fold dilution of goat anti-human kappa-HRP (SouthernBiotech, cat. no. 2060-05) was added to the plate and incubated for one hour at room temperature with gentle shaking. The plate was washed three times with PBST and developed with TMB substrate (Thermo Scientific, cat. no. 34028). The reaction was stopped with 2 M H_2_SO_4_. Absorbance was measured at 450 nm.

### Calcium flux assays

Fluo-4 AM was purchased from Thermo Fisher Scientific (cat. no. F14201). Fluo-4 AM loading solution (3 μM fluo-4 AM and 0.1% Pluronic F-127 in Hanks’ balanced salt solution (HBSS)) was prepared according to the manufacturer’s protocol. TMD8 cells (1 x 10^6 cells/mL) were incubated with Fluo-4 AM loading solution for 30 minutes at 37 °C and protected from light. Cells were washed once with complete medium and resuspended in complete medium at 1 x 10^6 cells/mL. Before each run, 500 μL of cell solution (1 x 10^6 cells/mL) was transferred to an Eppendorf tube and placed in a 37 °C heat block for exactly 15 minutes. Cells were then transferred to a 5 mL polystyrene tube, and baseline fluorescence was measured for one minute. After baseline, monoclonal anti-human IgM (αIgM) antibody was quickly added to the TMD8 cells (final concentration: 20 µg/mL). The cells were briefly vortexed and returned to the flow cytometer. Calcium mobilization was measured for five minutes. Fluorescent signal was recorded on a BD LSRFortessa Cell Analyzer.

### Quantitative analysis of FCRL5 and FcγRIIb expression on TMD8 cells

The expression levels of FCRL5 and FcγRIIb on TMD8 cells were quantified using the Quantum Simply Cellular (QSC) kit by Bangs Laboratories, Inc. (cat. no. 815A). The QSC kit includes five bead populations: one blank and four with increasing levels of Fc-specific capture antibody. The antibody binding capacity (ABC) of each bead population is pre- determined by the manufacturer. When the TMD8 cells and the five bead populations are stained with the same antibody, the ABC value can be determined for the stained TMD8 cells. The assay was carried out according to the manufacturer’s protocol. TMD8 cells and bead populations were stained using anti-FCRL5-PE mAb (BioLegend, cat. no. 340304), anti- FcγRIIb-APC mAb (BioLegend, cat. no. 398303), and corresponding isotype controls. Fluorescent signal was recorded on a BD LSRFortessa Cell Analyzer.

### Crystallization

Sitting-drop vapor diffusion method was employed to obtain diffraction-quality crystals. For crystallization of the rFCRL5 D1-D3:IgG1 Fc-K320E/Q342A complex, 1 µL of protein at 8.5 mg/mL and reservoir solution containing 0.1 M MES pH 6.5, 25% (w/v) PEG 8000 were mixed at 1:1 ratio in the HR3-159 Cryschem Plate (Hampton Research) at 22 °C. The final crystals for the complex were obtained after 5-7 days. Individual crystals were flash-frozen directly in liquid nitrogen after brief incubation with a reservoir solution supplemented with 30% (v/v) glycerol.

### Data collection and structure determination

X- ray diffraction data were collected at 24-ID-E beamline in Advanced Photon Source (Argonne National Lab). Complex data was collected using Eiger 16M detector at a wavelength of 0.9790 Å. All datasets were processed using XIA2 pipeline which is integrated in 24-ID-E auto processing platform. X-ray diffraction patterns were processed to 3.42 Å (FCRL5-Fc complex). In Phenix software, phases of FCRL5 and IgG1 Fc were obtained by molecular replacement using human IgG1 Fc (PDB: 5JII) and predicted FCRL5 (AlphaFold 3 (*32*)) as search model. Phase information was derived from the protein cores, excluding carbohydrates, and the resulting electron density maps revealed clearly distinguishable density at the expected locations of the glycosylation sites. The saccharide chains were manually built in Coot. All structures were iteratively built using Coot and Phenix refinement package. The quality of the finalized crystal structures was evaluated by MolProbity. The final statistics for data collection and structural determination are shown in Supplementary Table 1.

### Native mass spectrometry

rFCRL5 D1-D3 and trastuzumab-K320E/Q342A were individually buffer-exchanged into 100 mM ammonium bicarbonate at a concentration of 10 μM using Micro Bio-Spin P6 gel columns according to the manufacturer’s instructions. Following buffer exchange, the two proteins were mixed at equimolar concentrations (1:1 molar ratio) and incubated for 1 hour at room temperature to allow complex formation prior to native MS analysis. Approximately 2 μL of the resulting solution was loaded into gold/palladium-coated borosilicate static emitters pulled in-house and introduced by electrospray ionization using an applied voltage of 0.9 kV. Native mass spectra were acquired in the positive ion mode on a Q Exactive Plus UHMR mass spectrometer over an m/z range of 350–10,000 using a resolution setting of 1K at m/z 400. An in-source trapping voltage of −50 to −100 V was applied to enhance desolvation and transmission of intact non-covalent complexes.

The mass spectra obtained under native conditions were deconvoluted using UniDec (*63*). Default deconvolution parameters were applied with minor adjustments to accommodate the expected mass range of the analytes. Specifically, the mass range was set to 10–200 kDa and the charge state range to 1–35. The peak detection threshold was adjusted to 0.05–0.1, and the peak detection window was defined as 100–500 Da. Mass sampling was performed at 1 Da intervals. Peaks were fitted using a Gaussian function, and the charge smoothing width was set to 1.

## Acknowledgements

We thank Ed Satterwhite and Rocio Zapata Bustos for their help with protein expression. This work was supported by grants NIH U01AI148118, R35GM148356 to Y.J.Z., and R35GM139658 to J.S.B. Additionally, the Welch Foundation (F-1155) to J.S.B., Clayton Foundation to G.G., and the L. Leon Campbell Professorship are gratefully acknowledged. X-ray diffraction data collection was performed with support from an APS beam time award(s) (DOI: https://doi.org/10.46936/APS-191584/60015284) from the Advanced Photon Source, a U.S. Department of Energy (DOE) Office of Science user facility operated for the DOE Office of Science by Argonne National Laboratory under Contract No. DE-AC02- 06CH11357.

