## Supplementary Materials for "Structure-Function Analysis of the FCRL5-IgG1 Fc Complex Reveals an Unappreciated Effect of Fc-Silent Antibodies on B cells"

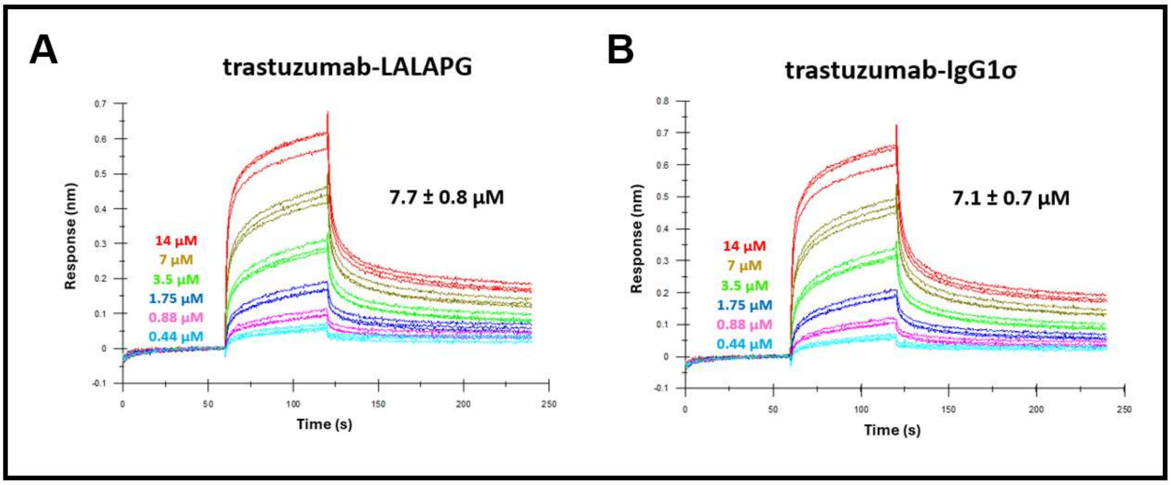


**Figure S1. BLI sensorgrams of Fc-silenced trastuzumab variants binding to FCRL5 (Q16-G851).** Antibodies were 2-fold serially diluted in kinetics buffer (PBS, 1% BSA, 0.02% Tween 20) starting at 14 μM. Triplicate measurements were performed for each antibody. The dissociation constant ($K_D$) is shown for each antibody as mean ± SD.

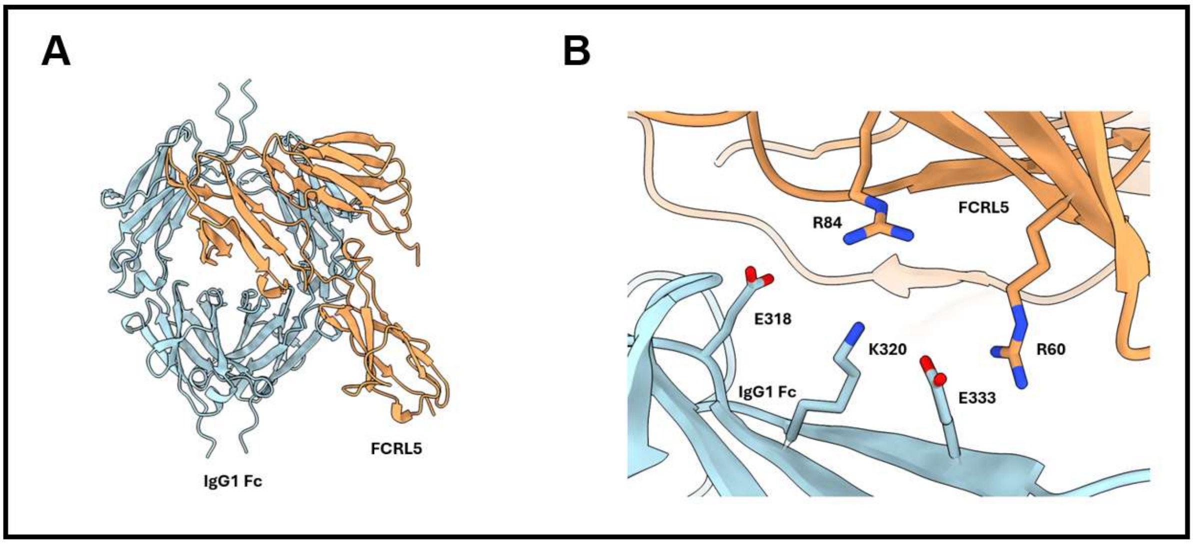


**Figure S2. FCRL5 D1-D3:IgG1 Fc complex structural prediction by AlphaFold 3.** (A) Overall predicted structure of the FCRL5 D1-D3 (Q16-I282):IgG1 Fc (EU numbering: D221-K447) complex. FCRL5 is shown in orange and IgG1 Fc in light blue. (B) Zoom in on the structural model for FCRL5 residues R60 and R84 and IgG1 Fc residues E318, K320, and E333. FCRL5 is shown in orange and IgG1 Fc in light blue.

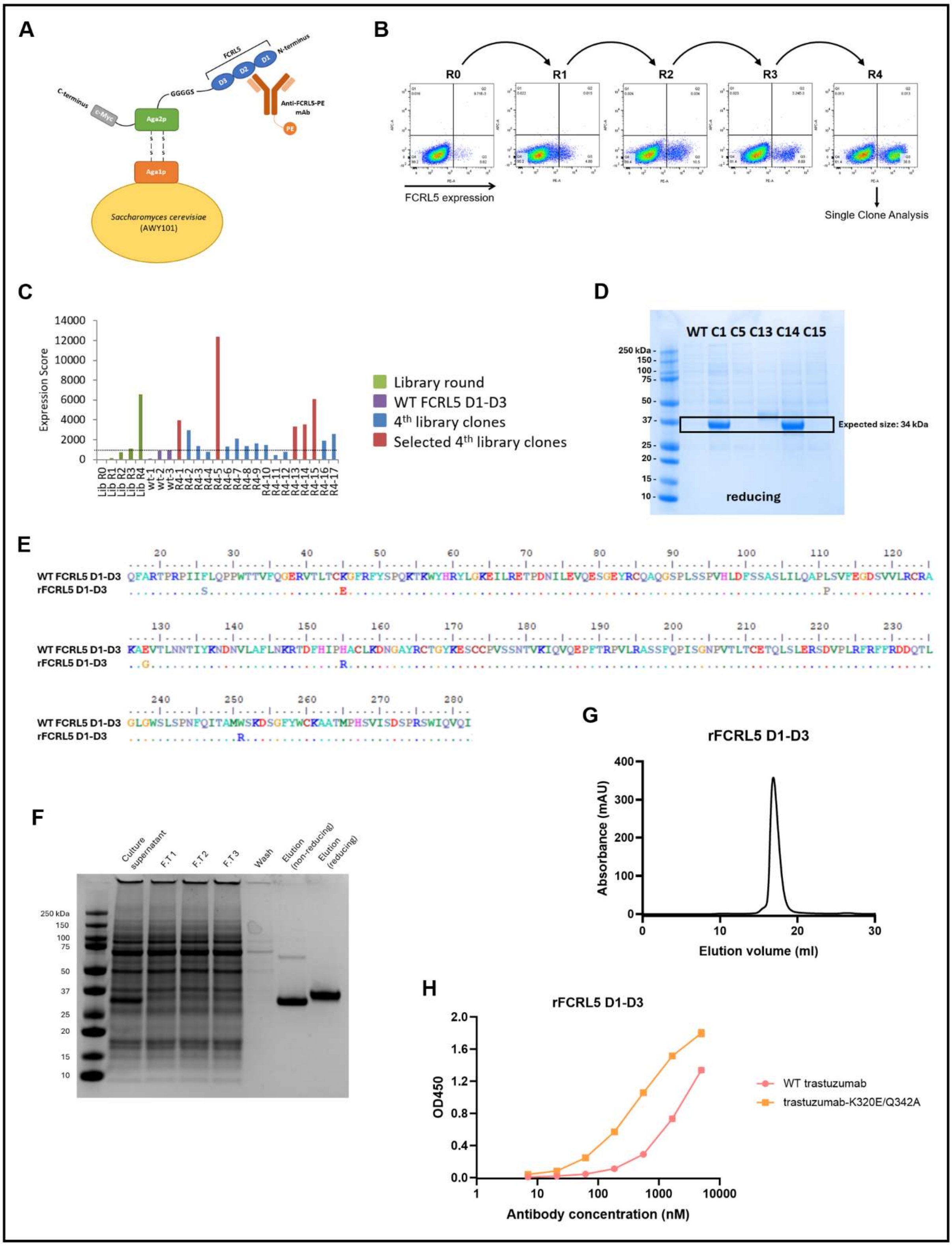


**Figure S3. Engineering of recombinant expression optimized rFCRL5 D1-D3.** (A) Schematic of Aga2p-FCRL5 D1–D3 on yeast. (B) Enrichment of FCRL5-expressing clones during sequential rounds of FACS. R0 represents the initial library, while R4 indicates the final sort. (C) Single clone analysis from round 4. The expression score was determined for each of the library rounds (R0-R4), three WT FCRL5 D1-D3 controls, and seventeen mutant clones from R4 (n = 1). The expression score was calculated as the percentage of cells in the high-expression quadrant (Q3) multiplied by the median fluorescence intensity (MFI) of that quadrant. The dotted line indicates the expression score of the WT controls. (D) Reducing SDS-PAGE showing the relative expression levels in Expi293F cells of WT FCRL5 D1-D3 and of the five clones from round 4 having the highest expression score (C1, C5, C13, C14, and C15). (E) Sequence alignment of the WT FCRL5 D1-D3 protein sequence with the highly expressed variant, rFCRL5 D1-D3. (F) SDS-PAGE of the different fractions collected during Ni-NTA affinity purification of rFCRL5 D1-D3. (G) SEC elution profile of purified rFCRL5 D1-D3. (H) ELISA measurement of rFCRL5 D1-D3 binding to WT trastuzumab and trastuzumab-K320E/Q342A.

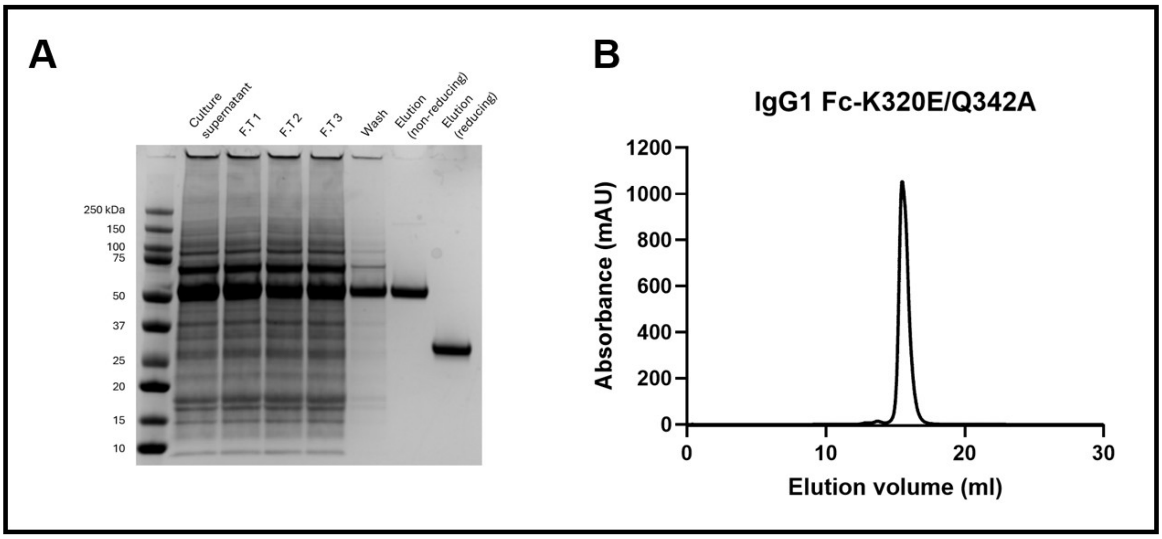


**Figure S4. Purification results for IgG1 Fc-K320E/Q342A**. (A) SDS-PAGE of the different fractions collected during protein G affinity purification of IgG1 Fc-K320E/Q342A. B) SEC elution profile for purified IgG1 Fc-K320E/Q342A.

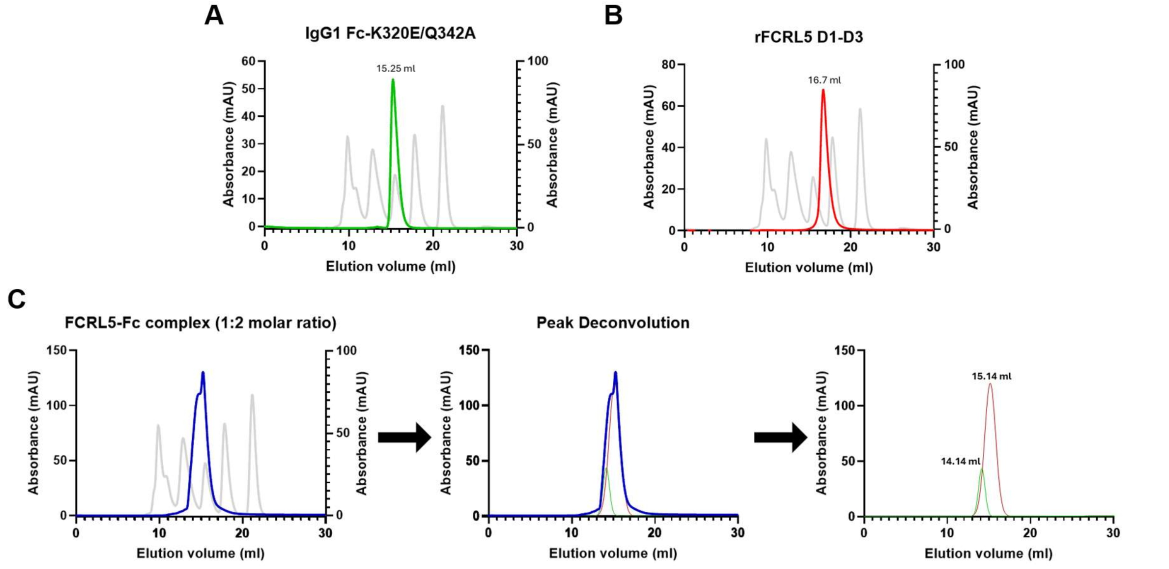


D

| Sample | Theoretical M.W. (kDa) | 1st peak eluted at (ml) | 2nd peak eluted at (ml) | 1st peak apparent M.W. (kDa) | 2nd peak apparent M.W. (kDa) |
|---|---|---|---|---|---|
| IgG1 Fc-K320E/Q342A | 50 | 15.25 | X | 45 | X |
| rFCRL5 D1-D3 | 34 | 16.7 | X | 21 | X |
| FCRL5-Fc complex | 84 | 14.14 | 15.14 | 83 | 48 |

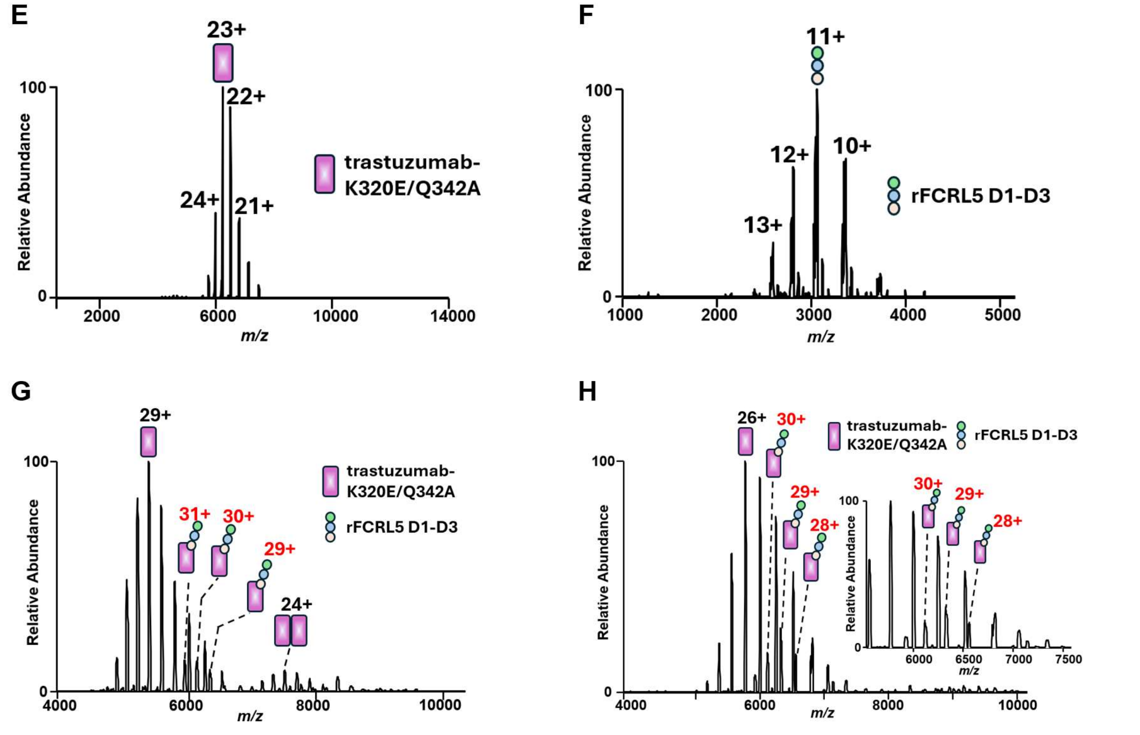


**Figure S5: Determination of the stoichiometry of the FCRL5-IgG complex.** (A-C) SEC elution profiles of (A) 28 μM IgG1 Fc-K320E/Q342A in PBS, (B) 28 μM rFCRL5 D1-D3 in PBS, and (C) 28 μM rFCRL5 D1-D3 mixed with 56 μM IgG1 Fc-K320E/Q342A in PBS. The elution profiles are overlaid with a calibration standard for reference. SEC traces are shown as absorbance versus elution volume, with the absorbance of the protein sample plotted on the left y-axis and the absorbance of the calibration standard plotted on the right y-axis. The overlapping peaks were deconvoluted by OriginPro. (D) A summary of the theoretical molecular weight (M.W.) of each protein (complex) and the apparent M.W. of each peak. (E) Native mass spectrum of trastuzumab-K320E/Q342A in 100 mM ammonium bicarbonate solution. (F) Native mass spectrometry of rFCRL5 D1-D3 in 100 mM ammonium bicarbonate solution. (G) Mass spectrum of a protein mixture containing 5 μM rFCRL5 D1-D3 and 5 μM trastuzumab-K320E/Q342A (1:1 molar ratio). (H) Mass spectrum of a protein mixture containing 9.1 μM rFCRL5 D1-D3 and 0.9 μM trastuzumab-K320E/Q342A (10:1 molar ratio). The inset shows an expanded view of the indicated m/z region, highlighting the peaks corresponding to the 1:1 FCRL5-IgG complex. Products corresponding to a 2:1 FCRL5-IgG complex were not detected.

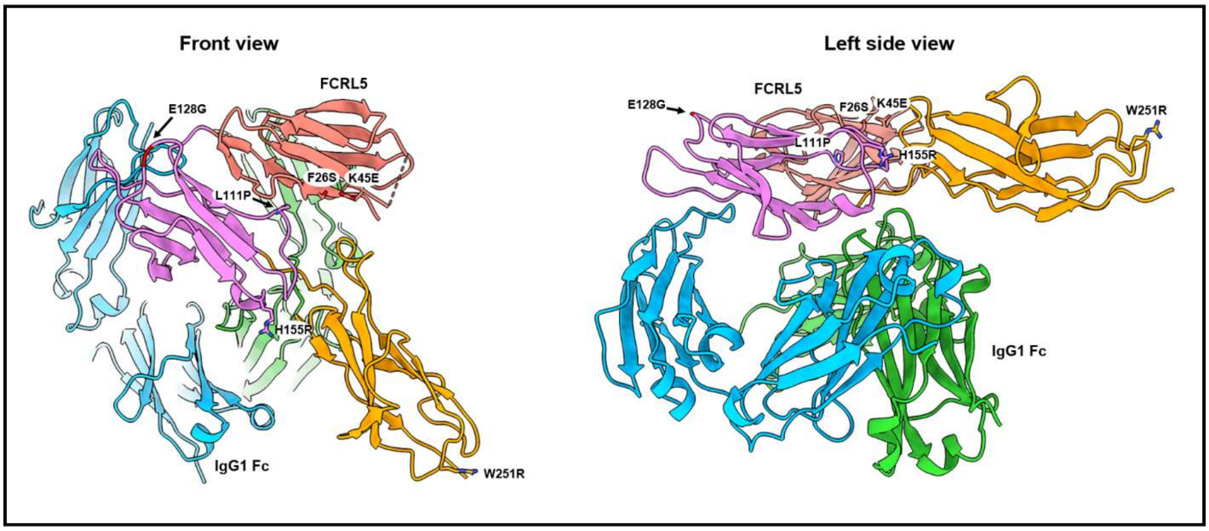


**Figure S6. The amino acid substitutions that enhance the expression of rFCRL5 D1-D3 are positioned away from the binding interface with IgG1 Fc.** Shown are a front view and a left side view of the crystal structure of rFCRL5 D1-D3 in complex with IgG1 Fc-K320E/Q342A. IgG1 Fc chains A and B are colored light blue and light green, respectively, while FCRL5 domains D1, D2, and D3 are shown in salmon, violet, and orange. The six substitutions in rFCRL5 D1-D3 (F26S, K45E, L111P, E128G, H155R, and W251R) are shown as sticks.

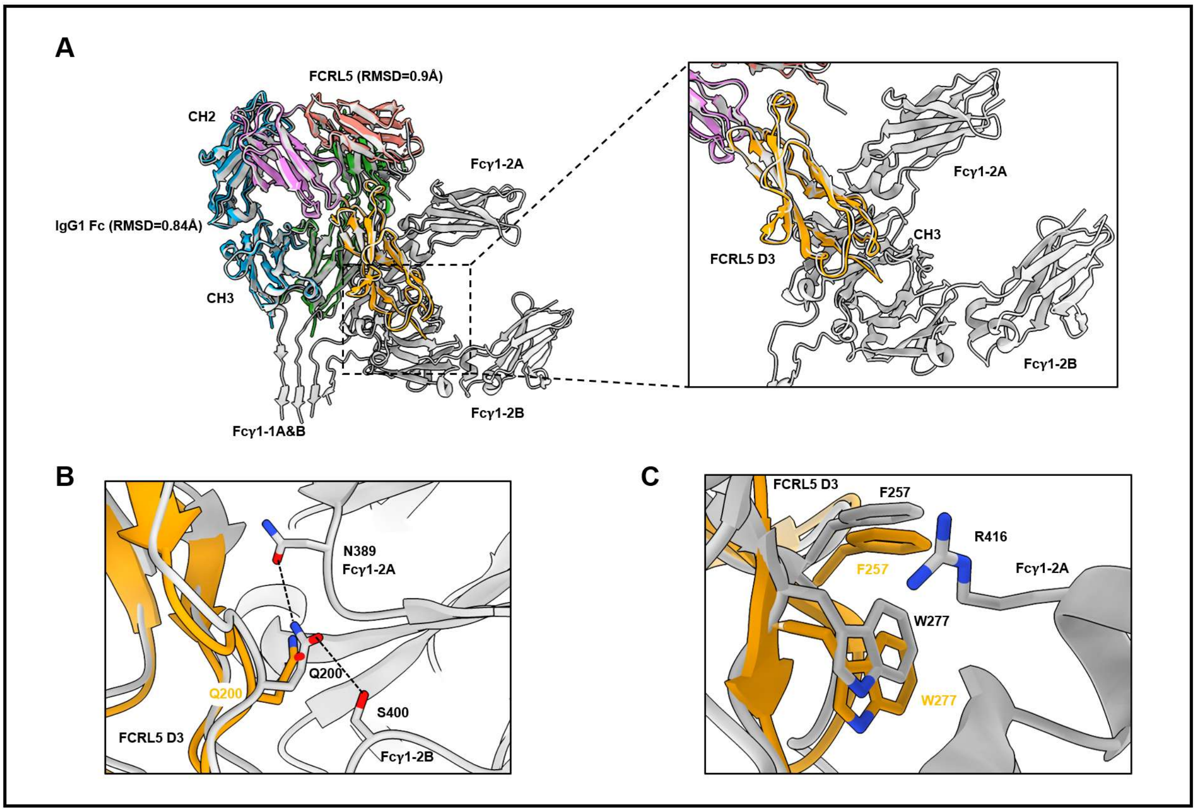


**Figure S7. Structural comparison of cryo-EM and crystal structures of the FCRL5–IgG1 Fc complex.** (A) Structural superposition of the cryo-EM FCRL5–IgG1 Fc complex (gray; PDB: 9LOC) with the crystal structure determined in this study (colored). The crystal structure is shown in the same colors as indicated in Figure 3A. The two structures align closely, with minimal conformational differences observed for both FCRL5 (Cα RMSD = 0.9 Å) and IgG1 Fc (Cα RMSD = 0.84 Å). The boxed region highlights the interface between the FCRL5 domain D3 and the CH3 domains of the second Fc molecule (Fcγ1-2). (B, C) Close-up views of the interface between FCRL5 D3 and Fcγ1-2. (B) Polar interactions observed in the cryo-EM structure. FCRL5 residue Q200 forms hydrogen bonds with Fcγ1-2 residues N389 and S400. (C) Close-up view of van der Waals interactions observed in the cryo-EM structure. Aromatic residues F257 and W277 of FCRL5 form van der Waals interactions with R416 of Fcγ1-2.

| | Crystal structure of human FCRL5 bound to IgG1 Fc |
|---|---|
| Wavelength (Å) | 0.9790 |
| Resolution range (Å) | 88.39 - 3.42 (3.54 - 3.42)$^{\pm}$ |
| Space group | P $3_1$21 |
| Unit cell: a, b, c (Å) | 138.72 138.72 130.51 |
| Unit cell: $\alpha, \beta, \gamma$ (°) | 90 90 120 |
| Total reflections | 414305 (43252) |
| Unique reflections | 20022 (1993) |
| Multiplicity | 20.7 (21.7) |
| Completeness (%) | 99.67 (99.05) |
| Mean I/sigma(I) | 3.57 (0.67) |
| Wilson B-factor(Å$^2$) | 88.78 |
| R-merge | 0.6715 (3.381) |
| R-pim | 0.1505 (0.7445) |
| $CC_{1/2}^{\Upsilon}$ | 0.957 (0.281) |
| CC* | 0.989 (0.662) |
| Refinement | |
| Reflections used in refinement | 19963 (1979) |
| Reflections used for R-free | 995 (93) |
| R-work | 0.2810 (0.3786) |
| R-free | 0.3060 (0.3529) |
| CC(work) | 0.884 (0.459) |
| CC(free) | 0.907 (0.445) |
| Number of non-hydrogen atoms | 5561 |
| Macromolecules | 5363 |
| Ligands | 198 |
| Protein residues | 674 |
| RMS(bonds) (Å) | 0.013 |
| RMS(angles) (°) | 1.63 |
| Ramachandran favored (%) | 98.04 |
| Ramachandran allowed (%) | 1.81 |
| Ramachandran outliers (%) | 0.15 |
| Rotamer outliers (%) | 0.32 |
| Average B-factor(Å$^2$) | 91.22 |
| Macromolecules | 89.98 |
| Ligands | 124.95 |
| Molprobity score^ | 1.75/100th percentile |

Statistics for the highest-resolution shell are shown in parentheses.

$^{\pm}$Values for the corresponding parameters in the outermost shell in parenthesis.

$^{\Upsilon}CC_{1/2}$ is the Pearson correlation coefficient for a random half of the data, the two numbers represent the lowest and highest resolution shell, respectively.

^MolProbity score is calculated by combining clashscore with rotamer and Ramachandran percentage and scaled based on X-ray resolution. The percentage is calculated with 100th percentile as the best and 0th percentile as the worst among structures of comparable resolution.

**Table S1. Data collection and refinement statistics.**
